## Supplementary figures and images for "Network Segregation Predicts Processing Speed in the Cognitively Healthy Oldest-old"

### Figure 1- supplemental figure 1

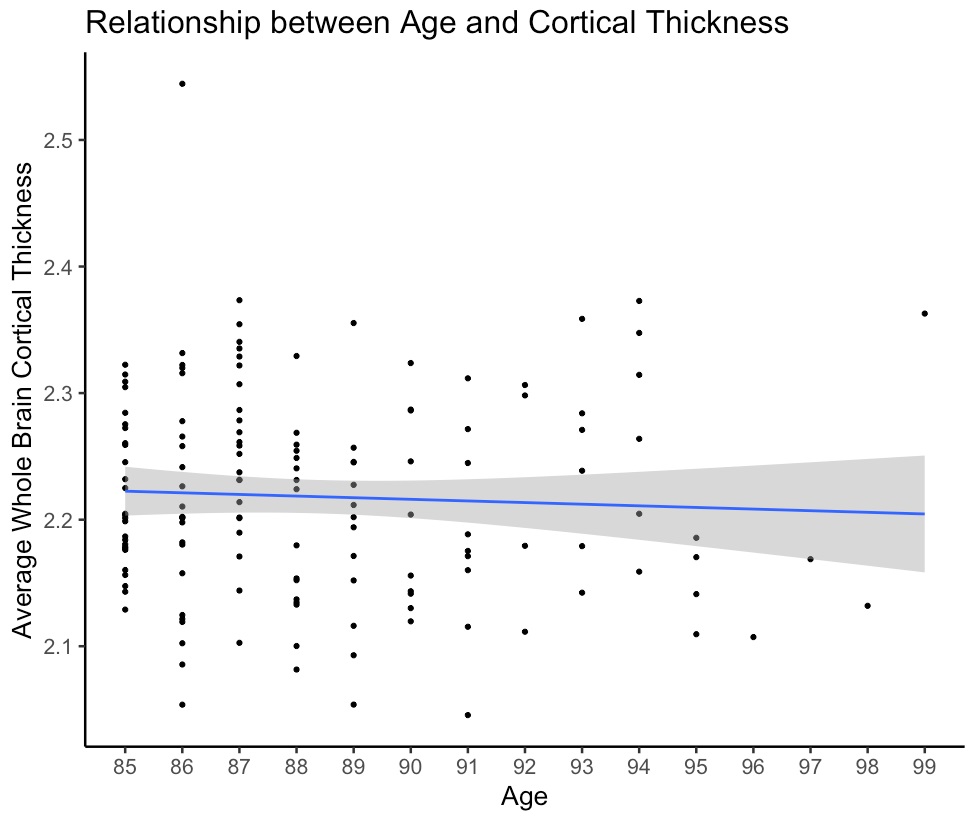

### Figure 6- supplemental figure 1

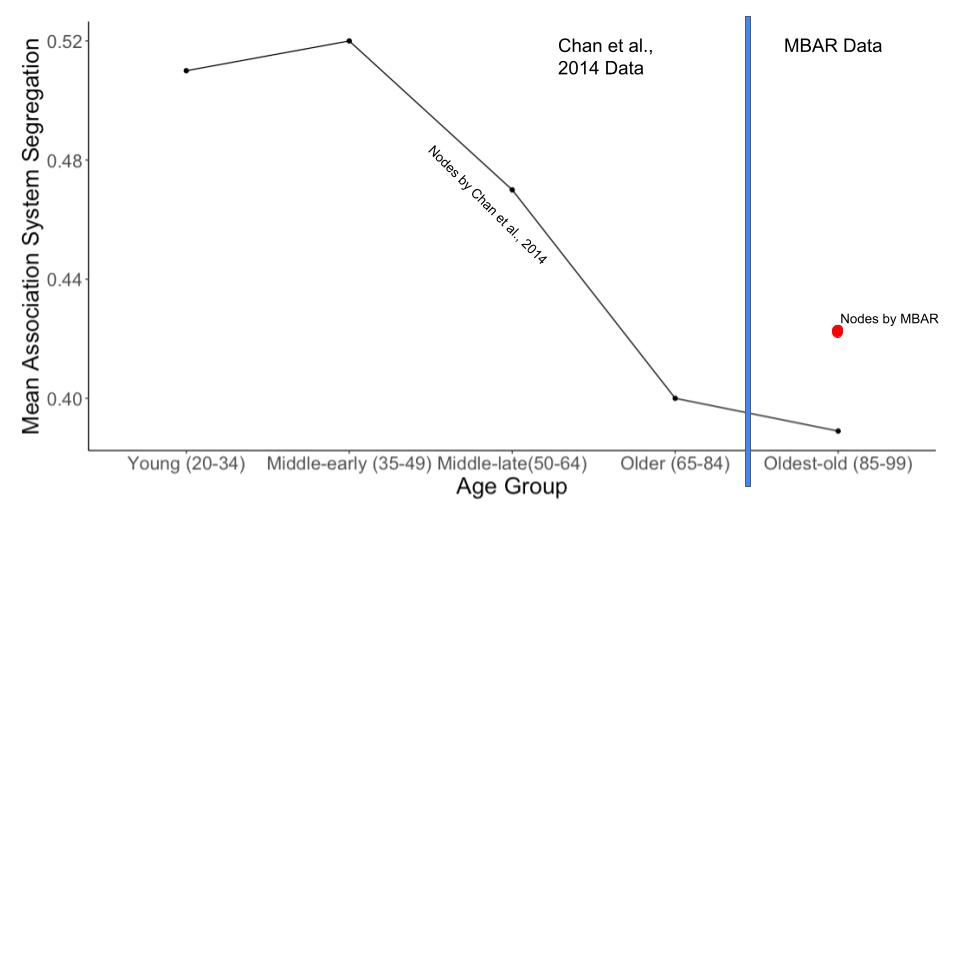

### Figure 7- supplemental figure 1

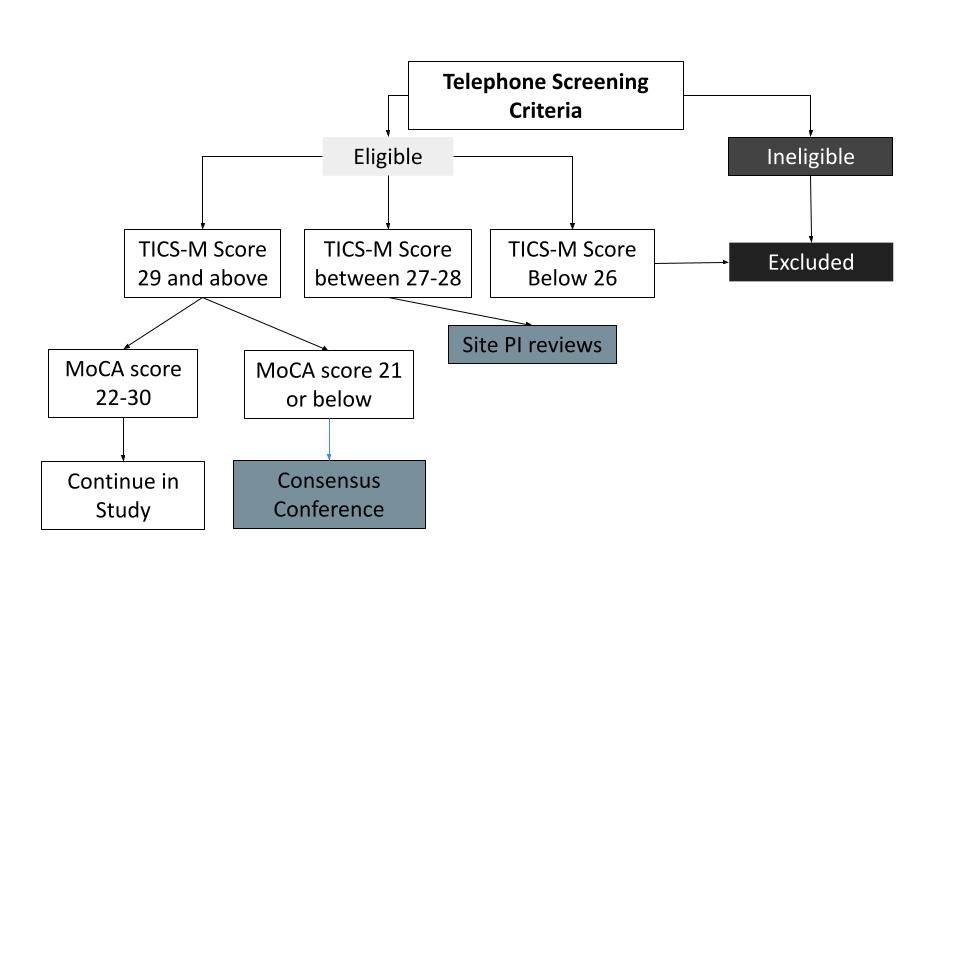
