## Supplementary File 1 for "Network Segregation Predicts Processing Speed in the Cognitively Healthy Oldest-old"

| **Participant Characteristics** | **Total, N=146** | **UAB^1^, N= 48 (32.9%)** | **UA^2^, N= 35 (24.0%)** | **UF^3^, N= 35 (24.0%)** | **UM^4^, N= 28 (19.2%)** |
| --- | --- | --- | --- | --- | --- |
| Age (years), *mean ± SD (range)* | 88.4 ± 3.18 (85-99) | 88.4 ± 3.47 (85-98) | 88.7 ± 2.98 (85-95) | 89.1 ± 3.65 (85-99) | 87.1 ± 1.70 (85-91) |
| Education (years), *mean ± SD (range)* | 16.1 ± 3.03 (9-26) | 15.7 ± 2.62 (12-22) | 15.9 ± 2.89 (9-22) | 16.5 ± 3.30 (10-22) | 16.5 ± 3.52 (12-26) |
| *Sex, N(%)* |  |  |  |  |  |
| Female | 79 (54.11%) | 24 (50.00%) | 18 (51.43%) | 20 (57.14%) | 17 (60.71%) |
| Male | 67 (45.89%) | 24 (50.00%) | 17 (48.57%) | 15 (42.86%) | 11 (39.29%) |
| *Race, N(%)* |  |  |  |  |  |
| Non-Hispanic Caucasian | 134 (91.78%) | 44 (91.67%) | 33 (94.29%) | 35 (100.00%) | 22 (78.57%) |
| African American | 6 (4.11%) | 4 (8.33%) | 0 (0.00%) | 0 (0.00%) | 2 (7.14%) |
| Hispanic Caucasian | 5 (3.42%) | 0 (0.00%) | 2 (5.71%) | 0 (0.00%) | 3 (10.71%) |
| Asian | 1 (0.69%) | 0 (0.00%) | 0 (0.00%) | 0 (0.00%) | 1 (3.57%) |
| *Marital Status, N(%)* |  |  |  |  |  |
| Widowed | 74 (50.69%) | 27 (56.25%) | 16 (45.71%) | 19 (54.29%) | 12 (42.86%) |
| Married | 54 (36.99%) | 17 (35.42%) | 12 (34.29%) | 13 (37.14%) | 12 (42.86%) |
| Divorced | 13 (8.90%) | 4 (8.33%) | 3 (8.57%) | 3 (8.57%) | 3 (10.71%) |
| Living as Married/Domestic Partnership | 3 (2.06%) | 0 (0.00%) | 3 (8.57%) | 0 (0.00%) | 0 (0.00%) |
| Never Married | 2 (1.37%) | 0 (0.00%) | 1 (2.86%) | 0 (0.00%) | 1 (3.57%) |
| *Dominant Hand, N(%)* |  |  |  |  |  |
| Right | 131 (89.73%) | 45 (93.75%) | 29 (82.86%) | 33 (94.29%) | 24 (85.71%) |
| Left | 15 (10.27%) | 3 (6.25%) | 6 (17.14%) | 2 (5.71%) | 4 (14.29%) |

**Participant Characteristics**: ^1^UAB, University of Alabama at Birmingham; ^2^UA, University of Arizona; ^3^UF, University of Florida; ^4^UM, University of Miami
