## Supplementary File 5 for "Network Segregation Predicts Processing Speed in the Cognitively Healthy Oldest-old"

| DV | **df** | **F** | **p-value** | **R^2^** |
| --- | --- | --- | --- | --- |
| *Overall Cognition* | 4 | 3.131 | 0.016 | 0.82 |
|  | **Standardized coefficient (β)** | **p-value** |  |  |
| *Association System Segregation* | 0.209 | 0.074 |  |  |
| *Association System Modularity* | 0.11 | 0.294 |  |  |
| *Association System Mean Connectivity* | 0.115 | 0.202 |  |  |
| *Association System Participation coefficient* | 0.098 | 0.355 |  |  |
| DV | **df** | **F** | **p-value** | **R^2^** |
| *Overall Cognition* | 3 | 6.026 | <.001 | 0.112 |
|  | **Standardized coefficient (β)** | **p-value** |  |  |
| *FPN Segregation* | 0.167 | 0.068 |  |  |
| *CON Segregation* | 0.139 | 0.139 |  |  |
| *DMN Segregation* | 0.115 | 0.259 |  |  |
| DV | **df** | **F** | **p-value** | **R^2^** |
| *Processing Speed* | 3 | 4.087 | 0.008 | 0.079 |
|  | **Standardized coefficient (β)** | **p-value** |  |  |
| *FPN Segregation* | 0.227 | 0.016 |  |  |
| *CON Segregation* | -0.056 | 0.553 |  |  |
| *DMN Segregation* | 0.119 | 0.252 |  |  |

**Supplemental Regressions:** Results from 3 multiple linear regressions: (1) association system metrics as predictors of overall cognition, (2) network segregation predictors of overall cognition, and (3) network segregation predictors of processing speed.
