## Supplementary File 3 for "Network Segregation Predicts Processing Speed in the Cognitively Healthy Oldest-old"

| **Parcellation System/Network** | **r** | **p** | **r with site cov** | **p with site cov** | **r with CT cov** | **p with CT cov** |
| --- | --- | --- | --- | --- | --- | --- |
| MBAR power association system | 0.342 | <.001 | 0.365 | <.001 | 0.351 | <.001 |
| MBAR power DMN | 0.273 | <.001 | 0.269 | 0.001 | 0.269 | 0.001 |
| MBAR power FPN | 0.272 | <.001 | 0.313 | <.001 | 0.275 | <.001 |
| MBAR power CON | 0.258 | 0.002 | 0.28 | <.001 | 0.256 | 0.002 |
| MBAR power SM system | 0.107 | 0.195 | - | - | - | - |
| Chan association system | 0.294 | <.001 | 0.324 | <.001 | 0.298 | <.001 |
| Chan DMN | 0.263 | 0.001 | 0.268 | 0.001 | 0.26 | 0.002 |
| Chan FPN | 0.255 | 0.002 | 0.278 | <.001 | 0.255 | 0.002 |
| Chan CON | 0.278 | <.001 | 0.304 | <.001 | 0.274 | <.001 |
| Chan SM system | 0.03 | 0.723 | - | - | - | - |
| Han association system | 0.261 | 0.001 | 0.3 | <.001 | 0.265 | 0.001 |
| Han DMN | 0.257 | 0.003 | 0.244 | 0.003 | 0.278 | <.001 |
| Han FPN | 0.32 | <.001 | 0.37 | <.001 | 0.331 | <.001 |
| Han CON | 0.197 | 0.017 | 0.24 | 0.004 | 0.192 | 0.021 |
| Han SM system | -0.058 | 0.489 | - | - | - | - |
| MBAR CD association system | 0.315 | <.001 | 0.339 | <.001 | 0.321 | <.001 |
| MBAR CD DMN | 0.271 | <.001 | 0.278 | <.001 | 0.268 | 0.001 |
| MBAR CD FPN | 0.254 | 0.002 | 0.307 | <.001 | 0.265 | 0.001 |
| MBAR CD CON | 0.2 | 0.015 | 0.215 | 0.009 | 0.19 | 0.02 |
| MBAR CD SM system | 0.036 | 0.663 | - | - | - | - |

**Correlations between Processing Speed and Segregation in each Parcellation:** Results of Pearson correlation between processing speed cognitive domain factor score and segregation of the association system, DMN, FPN, CON and Sensory motor system. Additionally, partial correlation results with site of data collection (site cov) and cortical thickness (CT cov) are included).
