## Supplementary File 4 for "Network Segregation Predicts Processing Speed in the Cognitively Healthy Oldest-old"

|  | **Memory** | **Working Memory** | **Language** | **Executive Functioning** |
| --- | --- | --- | --- | --- |
| FPN Segregation | r=.073  p=.382 | r=.124  p=.135 | r=.06  p=.473 | r=.167  p=.044 |
| DMN Segregation | r=-.017  p=.84 | r=.092  p=.729 | r=.137  p=.098 | r=.108  p=.192 |
| CON Segregation | r=-.102  p=.218 | r=.008  p=.921 | r=-.013  p=.869 | r=.08  p=.337 |

**Correlations between Cognitive Domains and Network Segregation:** results of a Pearson correlation between domain factor scores for each cognitive domain vs segregation of the three networks.
