## Supplementary File 2 for "Network Segregation Predicts Processing Speed in the Cognitively Healthy Oldest-old"

|  | **Processing Speed** | **Memory** | **Executive Functioning** | **Working Memory** | **Language** |
| --- | --- | --- | --- | --- | --- |
| WAIS-IV Coding^1^ | .785 |  |  |  |  |
| Stroop Color-Word Reading Trial^2^ | .678 |  |  |  |  |
| Trail Making Test A (lines/sec)^3^ | -.670 |  |  |  |  |
| WAIS-IV Symbol Search^1^ | .593 |  | .364 |  |  |
| CVLT II Long Delay Recall^4^ |  | .749 |  |  |  |
| FNAME Total Score^5^ |  | .709 |  |  |  |
| Craft Story Paraphrase Delay Recall^6^ |  | .599 |  |  |  |
| WAIS-IV Block Design^1^ |  |  | .704 |  |  |
| WAIS-IV Matrix Reasoning^1^ |  |  | .552 |  |  |
| Benson Figure Test Delay Recall^6^ |  | .362 | .376 |  |  |
| Stroop Color Word-Inhibition Test Interference^2^ |  |  | .367 |  |  |
| Trail Making Test B (lines/sec) (minus Trail Making Test A (lines/sec))^3^ |  |  | -.328 |  |  |
| Digit Span Forward^6^ |  |  |  | .745 |  |
| Digit Span Backward^6^ |  |  |  | .714 |  |
| WAIS-IV Letter-Number Sequencing^1^ |  |  |  | .362 |  |
| Letter Verbal Fluency (F & L)^7^ |  |  |  |  | .661 |
| WAIS-IV Similarities^1^ |  |  |  |  | .430 |
| Semantic Fluency (Animals)^8^ |  | .409 |  |  | .421 |

**Factor Loadings for Cognitive Domains:** results of factor analysis with cognitive tests. Loadings above .3 are included. ^1^ [(Weschler, 2008)](https://sciwheel.com/work/citation?ids=11279046&pre=&suf=&sa=0); ^2^ [(MacLeod, 1992)](https://sciwheel.com/work/citation?ids=372173&pre=&suf=&sa=0); ^3^ [(Gaudino, Geisler, & Squires, 1995)](https://sciwheel.com/work/citation?ids=4614290&pre=&suf=&sa=0); ^4^ [(Delis, Kramer, Kaplan, & Ober, 1987)](https://sciwheel.com/work/citation?ids=8625136&pre=&suf=&sa=0); ^5^ [(Amariglio et al., 2012)](https://sciwheel.com/work/citation?ids=2717775&pre=&suf=&sa=0); ^6^ [(Beekly et al., 2007)](https://sciwheel.com/work/citation?ids=8678997&pre=&suf=&sa=0); ^7^ [(Newcombe, 1969)](https://sciwheel.com/work/citation?ids=12158353&pre=&suf=&sa=0); ^8^ [(Benton, 1968)](https://sciwheel.com/work/citation?ids=5274496&pre=&suf=&sa=0)
